# The antifungal and other bioactive properties of the volatilome of *Streptomyces scabiei*

**DOI:** 10.1101/2025.04.24.650436

**Authors:** Nudzejma Stulanovic, Djulia Bensaada, Loïc Belde, Delphine Adam, Marc Hanikenne, Jean-François Focant, Pierre-Hugues Stefanuto, Sébastien Rigali

## Abstract

Volatile compounds (VCs) produced by most host-associated bacteria remain largely unexplored despite their potential roles in suppressing microbial competitors and facilitating host colonization. This study investigated the volatilome of *Streptomyces scabiei* 87-22, the model species for causative agents of common scab in root and tuber crops, under culture conditions that completely inhibited fungal growth, including the phytopathogens *Alternaria solani* and *Gibberella zeae*. Bicameral assays confirmed that these inhibitory effects were at least partially mediated by VCs. Using GC × GC TOFMS, 36 VCs were unambiguously identified as products of *S*. *scabiei* 87-22 metabolic activity. These included mainly ketones and aromatic compounds (both benzene derivatives and heterocycles), along with smaller contributions from other chemical families including sulfur-containing compounds, nitriles, esters, terpenoids, an amide, and an aldehyde. A literature survey suggests that many of these VCs possess antibacterial, antifungal, anti-oomycete, nematocidal, and insecticidal effects, while the bioactivity of others remains speculative, having been identified only within complex volatile mixtures. Among those with known antifungal properties, dimethyl trisulfide, 2-heptanone, and creosol inhibited the growth of the fungal pathogens tested in this study. In addition, we reveal here that 3-penten-2-one is also a strong inhibitor of fungal growth. Remarkably, despite *S*. *scabiei* 87-22 being defined as a pathogen, some of its VCs were associated with plant growth promotion and defense stimulation. Overall, our work highlights the remarkable potential of *S*. *scabiei* 87-22 to produce VCs with diverse bioactivities including fungal pathogen inhibition and priming plant resistance.

**Importance:** This study reveals that *Streptomyces scabiei*, the bacterium causing common scab in root and tuber crops, produces a wide variety of volatile chemicals with surprising benefits. These natural compounds can inhibit growth of other harmful microbes, including fungal plant pathogens. Some of these chemicals are already known to fight pests and diseases, while others, like 3-penten-2-one, are newly discovered as potential antifungals. Even more unexpectedly, some of the identified compounds may help plants grow or boost their defenses. Combined with previous work, our findings challenge the idea that *S*. *scabiei* is purely harmful and suggests it might, under certain conditions, play a protective role in its environment. This work deepens our understanding of how microbes interact with each other and with plants, and could help shape more sustainable approaches to agriculture.

## Introduction

Microbial communities within the rhizosphere are highly complex, both in their composition and the diverse roles played by various organisms, as well as the intricate web of interactions that occur among them (1–3). In addition to direct host-microbe interactions with plants, microorganisms engage in a variety of microbe-microbe interactions that can be antagonistic (competition), cooperative (mutualism), or neutral (commensalism), spanning both intra- and inter-kingdom dynamics (2, 4). Even within a single type of interaction such as antagonism, microorganisms employ multiple strategies to compete (2). One such strategy involves the production of volatile compounds (VCs) which can inhibit the growth of neighboring or distant organisms through direct toxicity, environmental pH modification, or by increasing competitors’ susceptibility to antibiotics (5). VCs are small molecules (<300 Da) with low boiling point and high vapor pressure, thereby facilitating interactions over relatively long distances (>20 cm) (4). While the term ’volatile’ primarily refers to their presence in the gas phase, these compounds can also diffuse into the aqueous phase. Despite their low solubility and generally apolar nature, their rapid diffusion into aqueous environments is facilitated by the absence of a hydration sphere (4).

A key focus of bacterial volatilome research is understanding the roles and modes of action of these VCs. Even for well-characterized compounds like geosmin, details regarding their production, perception, and biological functions are still emerging (6). This knowledge gap is partly due to the fact that VC production levels is highly dependent on specific environmental conditions including nutrient availability, temperature, pH, oxygen levels, and interactions with neighboring organisms, which may act as allies, antagonists or prey (4). In response to these changing conditions, VCs are emitted as chemical signals serving a variety of purposes, from antagonism to growth promotion, and to adaptation to biotic and abiotic stress. These signals shape microbial community dynamics and influence colonized plants, ultimately determining whether organisms escape, persist, invade or defend their ecological niche (7, 8).

The increasing recognition of the antimicrobial properties of VCs has fueled interest in developing ‘green’ alternatives to synthetic pesticides, particularly due to their rapid degradation in nature. Some VCs such as dimethyl disulfide, are already used in agriculture to control crop diseases (9, 10). However, their effectiveness is often limited by the concentration applied, and sometimes their use can even lead to undesirable effects and even to negative outcomes (11). Consequently, the search for effective and sustainable volatile-based agrochemicals or medicaments continues, with a focus on optimizing concentrations and enhancing synergistic interactions to improve efficacy against a broader spectrum of pathogens.

Although *Streptomyce*s species are widely studied for their symbiotic relationships and beneficial effects of their specialized metabolites, an increasing number of *Streptomyces* species and strains are documented for their phytopathogenic properties (12). The primary model species for studying plant pathogenic *Streptomyces* is *Streptomyces scabiei* 87-22 (syn. *scabies*), the causative agent of the common scab (CS) disease, which affects a wide range of root and tuber crops including potato, beet, carrot, radish, turnip, and parsnip (13). The virulence factors which predominantly contribute to the development of lesions are the thaxtomin phytotoxins (14). While much of the research to date has focused on the production and regulation of thaxtomins, growing attention is being directed towards understanding the function of other compounds like coronafacoyl phytotoxins, concanamycins, siderophores and phytohormones, their regulatory mechanisms, and their roles in plant colonization and rhizosphere community (15).

Importantly, despite causing surface lesions on tubers varying in size, shape, and texture, *S. scabiei* 87-22 and related pathogenic species rarely cause severe damage that compromises plant viability. Recently, we showed that the model species *Streptomyces scabiei* 87-22 inhibits the growth of plant pathogens, through siderophore-mediated iron sequestration in response to commonly used nitrogen fertilizers (ammonium nitrate, ammonium sulfate, sodium nitrate, and urea) (16). As *S. scabiei* does not cause lethal damage to its hosts and can inhibit the growth of more severe plant pathogens, the hypothesis that this species may, under certain conditions, function as a beneficial endophytic microorganism is currently debated within the scientific community. Building on these findings regarding the antagonistic behavior of *S. scabiei* 87-22, we explored here the volatilome of *S*. *scabiei* 87-22 as a potential source of antifungal volatile compounds that may provide a competitive advantage within the soil microbiome, aiding niche colonization and potentially hindering the growth of highly damaging plant pathogens.

## Results

### Overall antifungal activity of *S*. *scabiei* 87-22 according to culture conditions

*S*. *scabiei* 87-22 was grown under fifteen different culture conditions in order to identify those able to fully inhibit the growth of various fungal and oomycete microorganisms, suggesting the possible production of toxic VCs. Complete growth inhibition of *Alternaria solani* and *Penicillium restrictum* NS1 was observed in five media: ISP1, ISP6, LB, TSA, and MHA (Figure 1). *Fusarium culmorum* exhibited full inhibition only when *S*. *scabiei* 87-22 was cultivated on LB medium, while *Gibberella zeae* showed almost total growth inhibition on ISP1, ISP6, and TSA media. In contrast, the oomycete *Pythium ultimum* was only partially inhibited with the strongest inhibitory effect (33%) observed on ISP6 medium (Figure S1).

**Figure 1.**
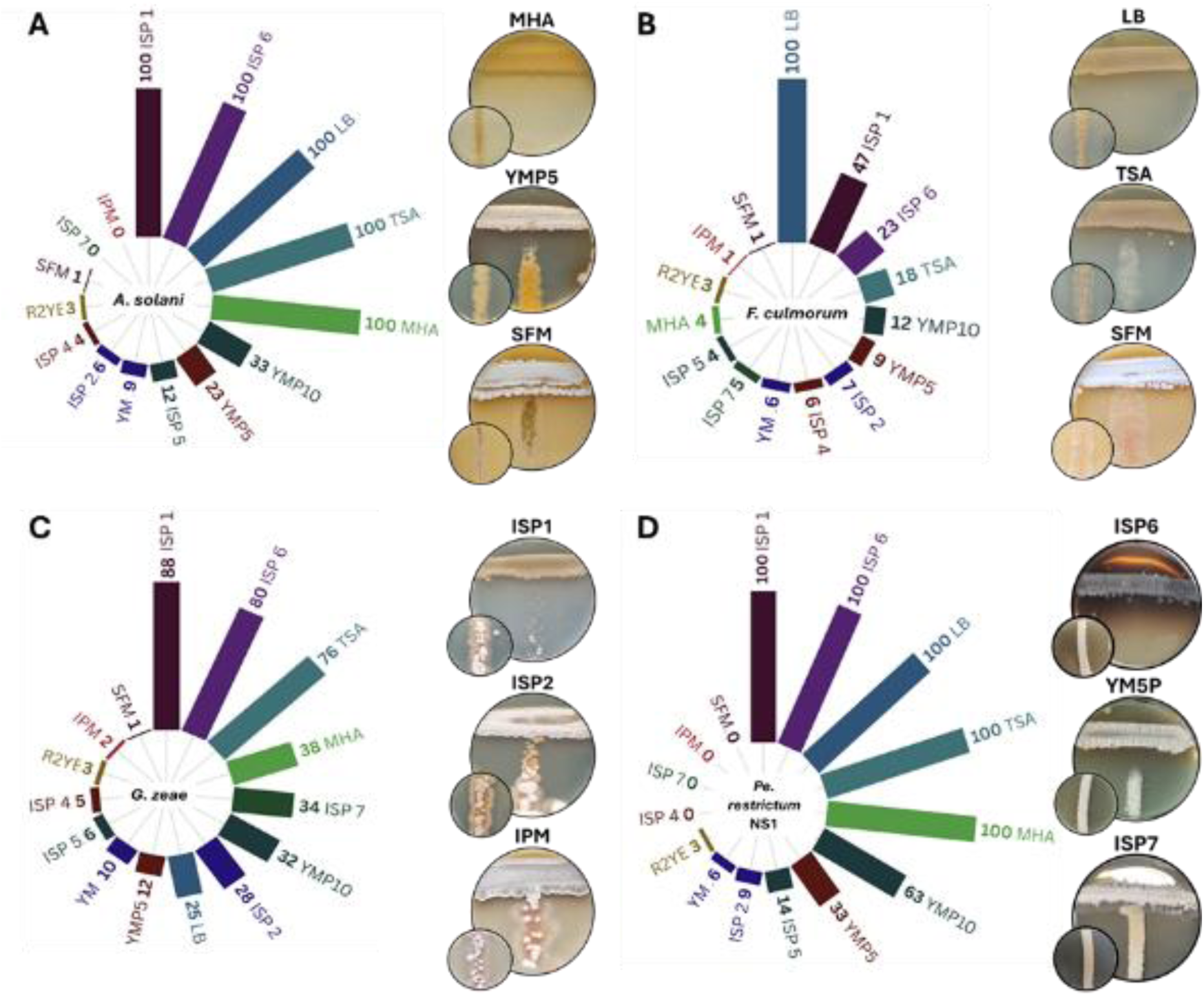
Semi-quantitative evaluation of antimicrobial activities of *S*. *scabiei* 87-22 according to the culture conditions. The bar plots display the inhibition scores from cross-streak assays with *S*. *scabiei* 87-22 grown on 15 different media. Tested microorganisms: (A) *Alternaria solani,* (B) *Fusarium culmorum,* (C) *Gibberella zeae,* (D) *Penicillium restrictum* NS1. A score of 100 means full growth inhibition of the tested microorganisms. Insets in the bottom left corner are control plates which display the growth of tested microorganisms without *S. scabiei.* Charts created using the Flourish studio (https://flourish.studio).

### *S*. *scabiei* 87-22 produces volatile compounds with antifungal properties

To examine whether these *S*. *scabiei* 87-22-mediated growth inhibition in cross-streak assays were caused by VC production, growth inhibition experiments were next conducted using bicameral Petri dishes. In these assays, dilution series of *Pe. restrictum* NS1 and streaks of *A. solani* and *G. zeae* were physically separated from a confluent lawn of *S. scabiei* 87-22 in the adjacent compartment (Figure 2). The VCs produced in ISP1, ISP6, LB, TSA, and MHA media resulted in complete inhibition of *Pe. restrictum* NS1, confirming that the antagonistic activity of *S*. *scabiei* 87-22 against this strain is at least partially mediated by VCs under these conditions (Figure 2). Growth inhibition assays against *A*. *solani* and *G*. *zeae* revealed varying degrees of inhibition depending on the medium used. Complete inhibition of both organisms was observed on ISP6 and TSA media, while *A*. *solani* was also entirely inhibited when *S*. *scabiei* 87-22 was grown on LB medium (Figure 2). On ISP1 and MHB media, both microorganisms also exhibited growth inhibition, though to a lesser extent compared to ISP6 and TSB (Figure 2). This partial growth inhibition of *A*. *solani* using bicameral Petri dishes (Figure 2B) suggests that the full growth inhibition observed in cross-streak assays on ISP1 and MHA (Figure 1A) necessitates both VCs and agar-diffusible antifungal compounds. Overall, *S*. *scabiei* 87-22 produces toxic VCs in all five media where maximal inhibition was previously observed in cross-streak assays (Figure 1).

**Figure 2.**
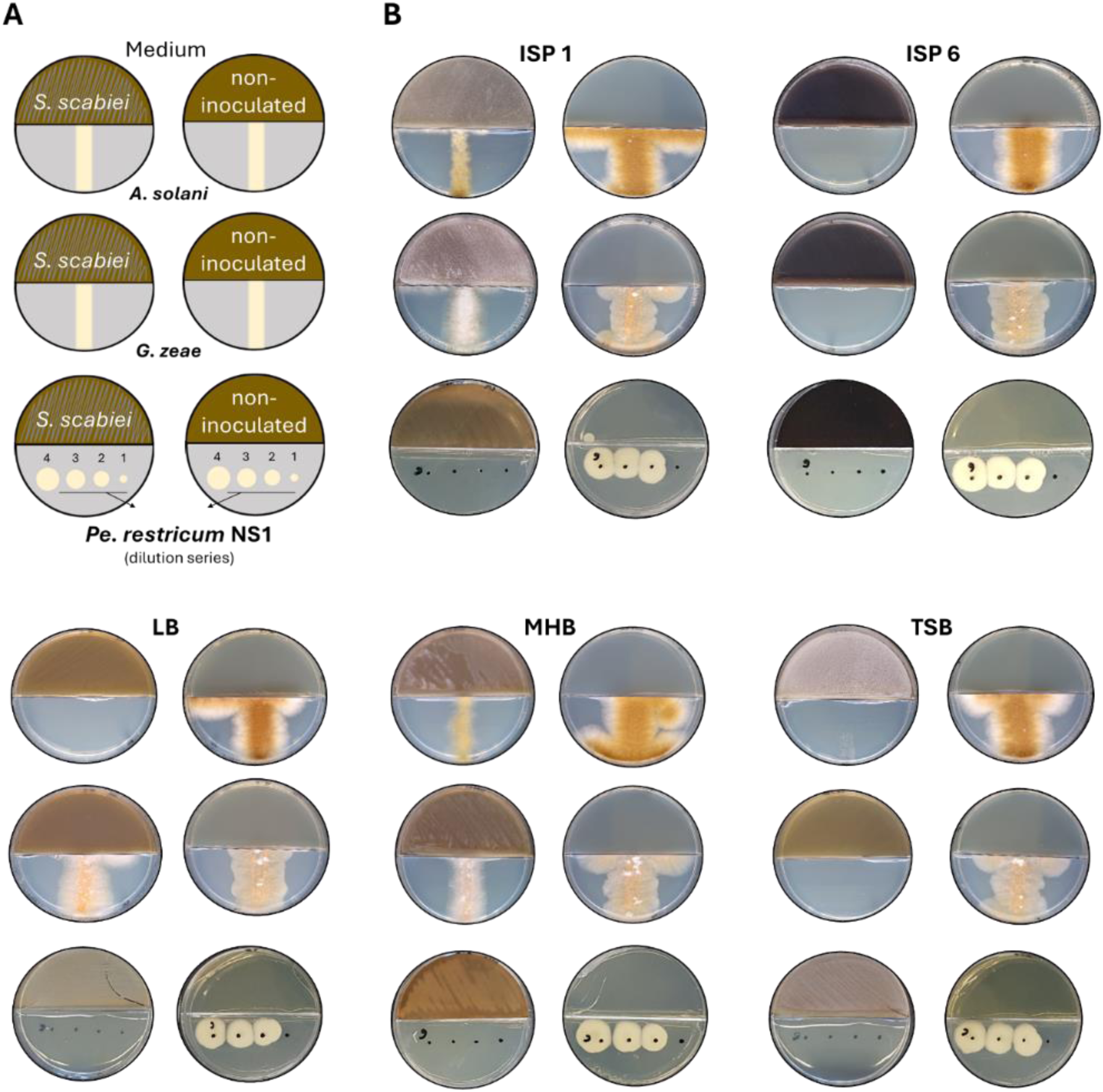
Evidence of antimicrobial activity of *S. scabiei* 87-22 mediated by VCs. **A.** Bicameral plate experiment setup**. B.** Assessment of volatile compounds of *S. scabiei* 87-22 active against *A. solani*, *G. zeae*, and *Pe. restrictum* NS1. The left plates with five different growth media (ISP1, ISP6, LB, MHB and TSB) were inoculated with *S. scabiei* 87-22, while the right plates served as growth controls (non-inoculated compartment).

Comprehensive two-dimensional gas chromatography coupled with time-of-flight mass spectrometry GC × GC–TOFMS was performed to identify VCs resulting from the metabolic activity of *S*. *scabiei* 87-22 grown under the five culture conditions (ISP1, ISP6, LB, TSA, and MHA) that trigger antimicrobial activity. Before conducting the comparative study, we confirmed that the VCs capable of inhibiting fungal growth were also produced in liquid ISP1, ISP6, LB, TSB (liquid TSA), and MHB (liquid MHA) (not shown). This step was crucial to facilitate compound extraction and enable direct sample injection for GC × GC–TOFMS analysis. The VCs in the culture supernatants of *S*. *scabiei* 87-22 were compared to those identified in these same media (non-inoculated controls), to identify compounds specifically resulting from the bacterium’s metabolic activity. The results, visualized as volcano plots (Figure S2), illustrate the volatilome of *S. scabiei* 87-22 across the five culture conditions tested (MHB, LB, ISP6, TSB, and ISP1). To ensure compliance with established standards, compound identification levels were assigned according to the guidelines set by the Metabolomics Standards Initiative (Supplementary Table S1) (17).

A Principal Component Analysis (PCA) revealed distinct VC fingerprints between the five non-inoculated media and their respective culture supernatant after growth of *S. scabiei* 87-22 (Figure 3A). The close grouping of replicates further reinforces the reliability and consistency of the measurements. The PCA results also suggest that the samples from the culture supernatants of LB, ISP6, and TSB occupy relatively independent spaces in the distribution map suggesting that the VCs produced are relatively different according to the culture medium composition. Interestingly, the VC composition of the culture supernatants of *S. scabiei* 87-22 grown in MHB and ISP1 showed similar profiles compared to the other three culture supernatants (Figure 3A). Their similar VC fingerprints may result from comparable nutrient composition of both media, containing casein hydrolysates (17.5 g/L in MHB and 5 g/L in ISP1) and either beef (2 g/L in MHB) or yeast (3 g/L in ISP1) extracts, with starch (1.5 g/L) as additional carbon sources in MHB. Despite these compositional overlaps and the similar VC profiles, the antimicrobial activities of *S*. *scabiei* differed between the two media.

**Figure 3.**
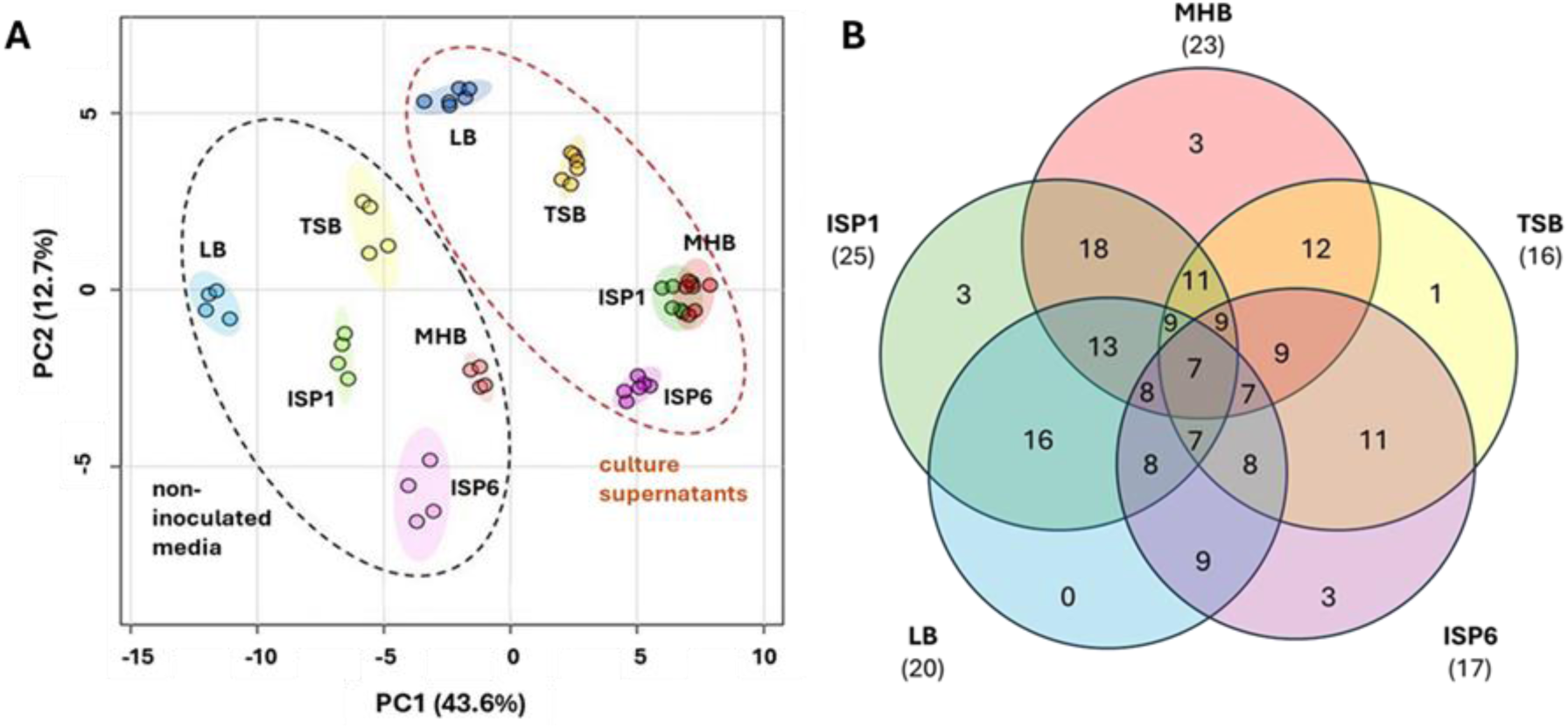
Volatilome profiling by gas mass spectrometry in *S. scabiei* 87-22 grown in five media. **A.** Principal component analysis (PCA) illustrating the distribution of volatile compounds. PC1 and PC2 account for 43.6% and 12.7% of the total variance, respectively, together accounting for 56.3% of the variability in the compound profiles and effectively distinguishing between non-inoculated media (black dotted ellipse) and culture supernatants (red dotted ellipse). (Biological replicates n = 2; technical replicates for inoculated media n = 3, technical replicates for non-inoculated media n = 2) **B.** The Venn diagram illustrates the number of VCs that are either specific to one medium or shared between two or more culture supernatants. The number below each medium name represents the total VCs with Log_2_ FC ≥ 1 detected in that medium.

The total number of detected VCs ranged from 200 to 400, depending on the culture medium. Of these, 36 were most likely derived from the metabolic activity of *S. scabiei* 87-22, exhibiting a significant increase in production (Log_2_ FC ≥ 1) compared to the control non-inoculated media (Table 1, Table S1 for MSI levels). Specifically, 25, 23, 20, 17 and 16 out of 36 VCs were significantly (Log_2_ FC ≥ 1) produced in ISP1, MHB, LB, ISP6, and TSB, respectively (Figure 3B,). Seven VCs were consistently produced across all five media (Figure 3B): 3-penten-2-one (F38), 2,2-dimethylheptane-3,5-dione (F65), dimethyl selenodisulfide (F35), 2-heptanone, 5-methyl (F25), 4-decanone (F68), benzoic acid methyl ester (F11), and 2-heptanone, 6-methyl (F37) (Table 1). 18 out of 36 VCs are shared in the ISP1 and MHB culture supernatants (Figure 3B), confirming their similar VC fingerprints as deduced from the PCA analysis. Figure 4A shows the structure of the 36 VCs identified in the volatilome of *S. scabiei*. The list of 36 volatile compounds covers several chemical families and reflects a broad chemical diversity. Ketones are the most abundant, accounting for 17 compounds in total (F25, F37, F38, F43, F46, F47, F50, F53, F56, F57, F64, F65, F68, F74, F76, F77, F79). Aromatic compounds are also well represented, comprising 10 compounds (F04, F11, F14, F17, F32, F36, F40, F66, F67, F71) that include both benzene derivatives and heterocycles. Sulfur-containing compounds follow, with four compounds (F19, F32, F35, F58) contributing to the diversity of functional groups. In addition, there are two nitriles (F02, F14), along with smaller contributions from other families, including esters (F40, F76) and terpenoids (F05, F06) and single representatives of an amide (F04), and an aldehyde (F60).

**Figure 4.**
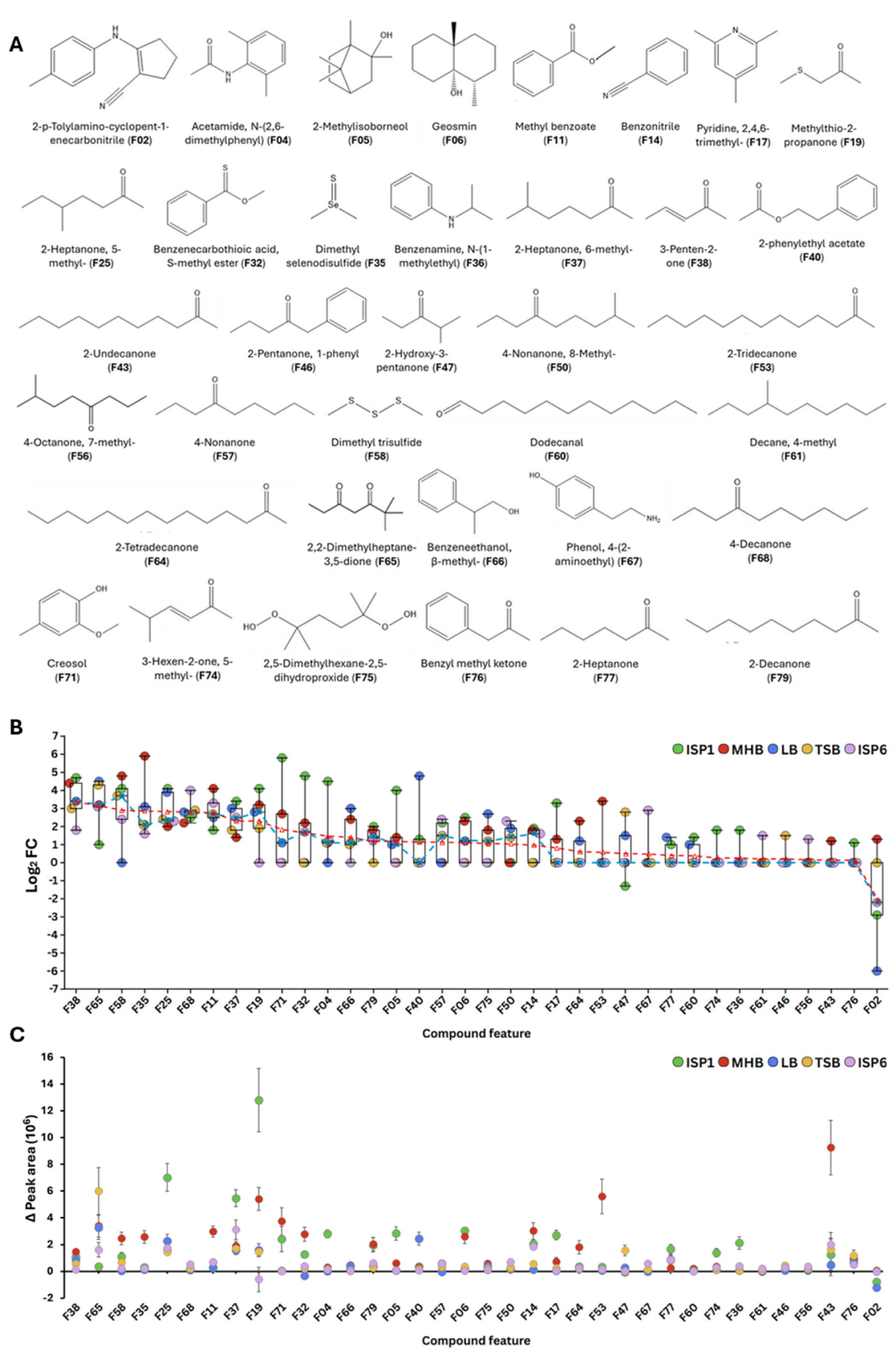
Structure, production and peak area variation of VCs produced by *S. scabiei* 87-22. **(A)**. Structures of the 36 identified VCs ranked by feature (F) numbers. (**B)** The 36 VCs ranked by their decreasing average Log_2_ FC value (presented by red dotted line) across the five culture conditions. The horizontal line within each box represents the median of five values, while the box itself spans from the first to the third quartile. The dotted blue line with cross represents the median Log_2_ FC of each VC across the five growth media. (**C).** Peak area variation of each VCs across the five media. Standard deviation of each dot shows the distribution of values derived from three technical replicates and two biological replicates. Biological replicates n = 2; technical replicates for inoculated media n = 3, technical replicates for non-inoculated media n = 2. Figure created with Flourish studio (https://flourish.studio).

**Table 1.** Compounds with significant Log_2_ FC ≥ 1 in at least one experimental condition and their reported activities.

| Feature number | Average Log<sub>2</sub> FC | Compound name | Reported activity | Ref |
| --- | --- | --- | --- | --- |
| **F58** | 2.99 | **Dimethyl trisulfide** | Antibacterial, Antifungal, Anti-oomycete, Nematicide | (18–21) |
| **F38** | 3.45 | **3-Penten-2-one** | NO synthase inhibitor | (22) |
| F65 | 3.24 | 2,2-Dimethylheptane-3,5-dione | Unknown | - |
| F19 | 2.39 | Methylthio-2-propanone | Unknown | - |
| F68 | 2.87 | 4-Decanone | Found in plant growth promoting extract | (23) |
| F11 | 2.86 | Methyl benzoate | Insect attractant and insecticide, Induced systemic resistance in plants | (24–26) |
| F37 | 2.39 | 2-Heptanone, 6-methyl- | Antifungal | (27, 28) |
| F25 | 2.92 | 2-Heptanone, 5-methyl- | Found in antifungal extracts | (21, 29) |
| F35 | 2.95 | Dimethyl selenodisulfide | Unknown | - |
| F32 | 1.74 | Benzenecarbothioic acid, S-methyl ester | Antifungal and antibacterial | (30) |
| F14 | 1.06 | Benzonitrile | Stimulates growth of *Pseudomonas fluorescens*, Found in antifungal extracts, Precursor for toxic and antimicrobial derivatives | (31–33) |
| F57 | 1.22 | 4-Nonanone | Unknown | - |
| F79 | 1.28 | 2-Decanone | Antifungal, Found in potato growth stimulating extracts | (21, 34) |
| F50 | 1.14 | 4-Nonanone, 8-Methyl- | Unknown | - |
| F06 | 1.21 | Geosmin | Antifungal, Insect attractant, Predation deterrent | (35–38) |
| F75 | 1.15 | 2,5-Dimethylhexane-2,5-dihydroperoxide | Unknown | - |
| **F71** | 1.92 | **Creosol** | Antifungal | (39) |
| F04 | 1.56 | Acetamide, N-(2,6-dimethylphenyl)- | Unknown | - |
| F66 | 1.50 | Benzeneethanol, β-methyl- | Unknown | - |
| F05 | 1.27 | 2-Methylisoborneol | Insect attractant, Predation deterrent | (36, 37) |
| F67 | 0.57 | Phenol, 4-(2-aminoethyl) | Neurotransmitter, Cholinesterase inhibitor | (40–42) |
| F56 | 0.25 | 4-Octanone, 7-methyl- | Found in insect repellent extracts | (43) |
| F61 | 0.30 | Decane, 4-methyl- | Found in antifungal and antibacterial extracts, Found in potato growth stimulating extracts | (44–46) |
| F46 | 0.30 | 2-Pentanone, 1-phenyl | Sexual signaling in arthropods | (47) |
| F47 | 0.58 | 2-Hydroxy-3-pentanone | Found in plant growth promoting extracts | (48, 49) |
| F53 | 0.67 | 2-Tridecanone | Found in antifungal extracts, Repellent, Nematicide<br>Found in potato growth stimulating extracts, Systemic Resistance induction in *Arabidopsis* | (21, 29, 46, 50–52) |
| F64 | 0.71 | 2-Tetradecanone | Antibacterial | (53) |
| F43 | 0.24 | 2-Undecanone | Antioomycete, Antifungal<br>Found in potato growth stimulating extracts | (46, 54, 55) |
| F17 | 0.91 | Pyridine, 2,4,6-trimethyl- | Antifungal | (56) |
| F60 | 0.47 | Dodecanal | Antifungal and antibacterial | (57) |
| F36 | 0.35 | Benzenamine, N-(1-methylethyl) | Unknown | - |
| F76 | 0.22 | Benzyl methyl ketone | Found in antifungal extracts | (21) |
| **F40** | 1.23 | **2-Phenylethyl acetate** | Antifungal | (58, 59) |
| **F77** | 0.49 | **2-Heptanone** | Antifungal, Antibacterial, Promote plant growth, Nematicide, Insecticide | (29, 31, 51, 60–62) |
| F74 | 0.36 | 3-Hexen-2-one, 5-methyl- | Unknown | - |
| F02 | -1.96 | 2-p-Tolylamino-cyclopent-1-enecarbonitrile | Unknown | - |

Figure 4B ranks these 36 VCs by decreasing average Log_2_ FC, highlighting consistent production of certain volatiles while others exhibit strong medium-dependent production variation. Nine compounds, including the seven VCs consistently produced (Log_2_ FC ≥ 1) across all five media (Figure 3B), showed a median Log_2_ FC above 2, indicating robust production across multiple media. The remaining two, dimethyl trisulfide (DMTS, F58) and methylthio-2-propanone (F19) were not produced in LB and TSB, respectively. For the other 27 compounds (median Log_2_ FC < 2), production varied widely across media, with some not produced in certain conditions while reaching peak production levels in others. For instance, creosol (F71) was undetectable in TSB and ISP6 but showed a high Log_2_ FC in ISP1 (5.81) and MHB (2.66), while 2-phenylethyl acetate (F40) was absent in most conditions but peaked at Log_2_ FC 4.84 in LB. Similarly, acetamide, N-(2,6-dimethylphenyl)-(F04), and 2-methylisoborneol (F05) were undetectable in some media yet exhibited significant overproduction in others (Figure 4B). A similar trend was observed when analyzing the average Log_2_ FC values across the five media (Figure 4B), reinforcing the median-based findings. Notably, compounds with high average Log_2_ FC often had relatively lower deviations, indicating stable production, whereas those with strong media dependence showed greater fluctuations, as reflected by higher standard deviations.

The comparison of the variation of peak areas of VCs across five different culture media (ISP1, MHB, LB, TSB, ISP6) provides valuable insights into the compounds that dominate the volatilome of *S*. *scabiei* 87-22 (Figure 4C). Certain compounds, such as F19 (methylthio-2-propanone), F25 (2-heptanone, 5-methyl-), F53 (2-tridecanone), and F43 (2-undecanone), exhibit high peak areas, exceptionally in ISP1 and MHB (Figure 4C). Notably, F19 and F25 show a sharp peak in ISP1, while F43 and F53 dominate in MHB. In contrast to MHB and ISP1 media, F19 exhibits a negative peak area variation in ISP6, suggesting that this compound is either not produced, consumed by *S. scabiei*, or it has served as a precursor for the production of another molecule. This observation aligns with a broader trend, as ISP6, along with LB and TSB, shows lower peak area tendencies across most compounds with only few exceptions in these media.

A literature survey was conducted on the 36 VCs to identify those with already known bioactivities, particularly antimicrobial properties, that could explain the observed growth inhibitions. The investigated VCs are associated with approximately ten distinct biological activities, including antimicrobial effects (antibacterial, antifungal, and anti-oomycete), as well as nematocidal and insecticidal properties (Figure 5, Table 1). More surprisingly for a species considered as a plant pathogen, some VCs positively impact plant development-related activities, including plant growth promotion and plant defenses. For some compounds, the biological activity has been confirmed through tests using the pure standards. For others, the reported activity remains speculative, as the compound was identified within a complex mixture of VCs, and its specific contribution to the overall effect has yet to be determined. Obviously, several VCs exhibit more than one activity (see for example F43, F53, F58, and F77 in Figure 5), while no bioactivity of 11 compounds could be retrieved from the literature (Figure 5, Table 1).

**Figure 5.**
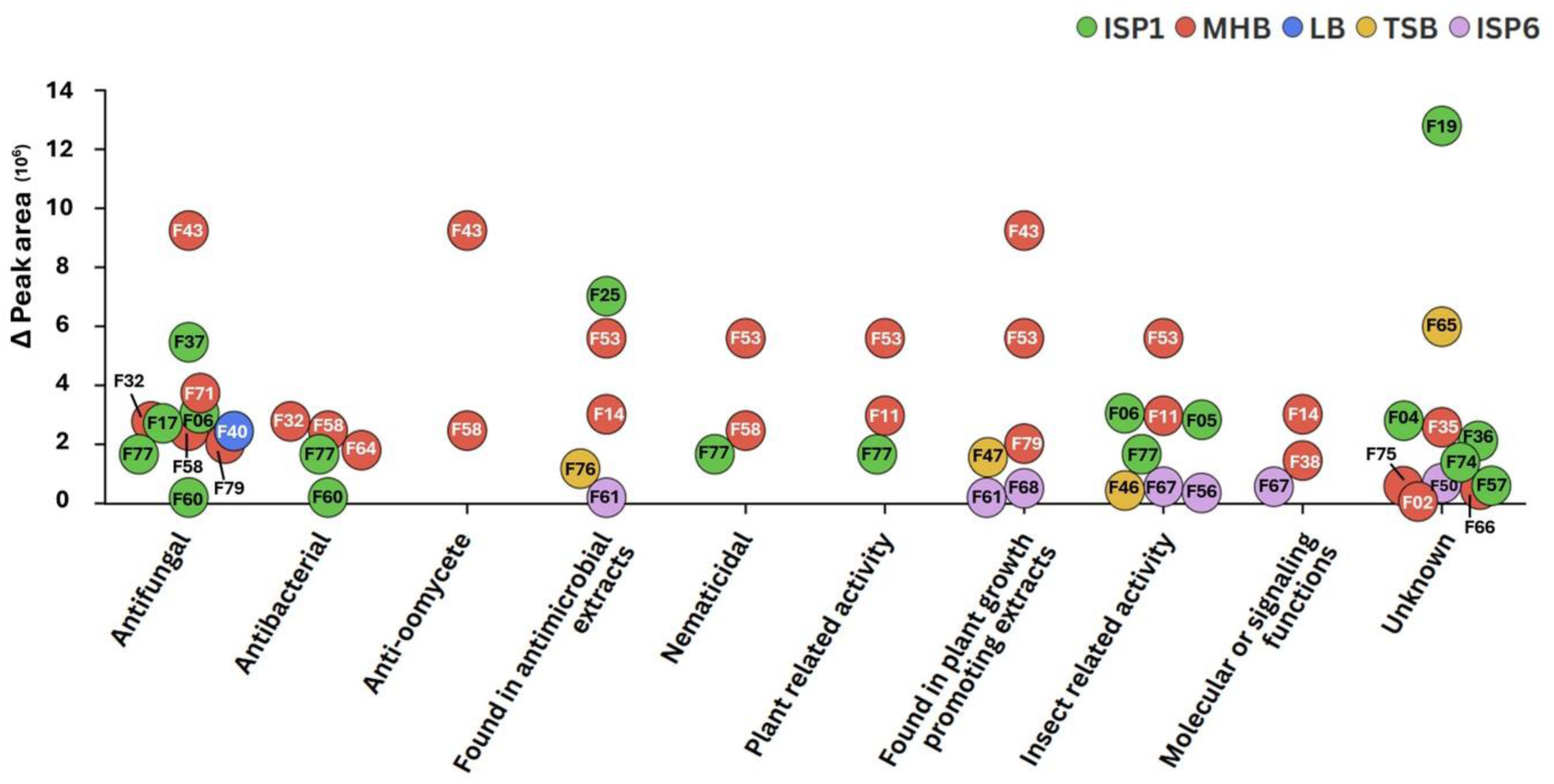
Functional distribution of volatile compounds produced by *Streptomyces scabiei* 87-22. Each dot represents the highest area value of an individual volatile compound, color-coded according to the culture medium in which it was most abundantly detected (ISP1, MHB, LB, TSB, ISP6). Compounds are labeled by their feature number (F), and their biological activities are categorized along the x-axis.

The candidate VCs responsible for the growth inhibition of *A*. *solani, G*. *zeae*, and *P*. *restrictum* NS1 have been first searched among the 12 VCs that have previously been reported for their antimicrobial properties (Table 1). **Dimethyl trisulfide** (DMTS) (F58) is well known for its broad antimicrobial activity against various bacteria (*Escherichia coli* O157:H7), pathogenic fungi (*Fusarium solani, Alternaria alternata*), and even oomycetes like *Phytophthora infestans*, the causative agent of late blight in potatoes (18–21, 63, 64). Other ketones, such as **2-heptanone** (F77), have been shown to suppress the growth of phytopathogenic fungi (*Fusarium oxysporum* (62) and *Moniliophthora roreri* (31)), phytopathogenic bacteria (*Agrobacterium tumefaciens* C58), nematodes, and insects (61), while positively influencing plant growth (60). **2-heptanone, 6-methyl** (F37) exhibited activity against *Alternaria solani* (27, 28). Similarly, **2-decanone** (F79) has demonstrated complete inhibition of *Fusarium oxysporum* (65) and *Fusarium solani* (21), as well as against *Colletotrichum acutatum* (66) and *Colletotrichum gloeosporioides* (67), both of which cause anthracnose in plants. Other compounds with strong antifungal activity include **2-phenylethyl acetate** (F40), which inhibited *Aspergillus ochraceus* (59), and **creosol** (F71), which completely suppressed *Fusarium*, *Penicillium*, and *Aspergillus* species (39). The inhibitory effects of **2-undecanone** (F43) were confirmed against the phytopathogenic fungus *Sclerotinia sclerotiorum* (54) and the phytopathogenic oomycete *Pythium myriotylum* (55). **Dodecanal** (F60) demonstrated antifungal activity against *Rhizoctonia solani* and antibacterial activity against the rice pathogen *Xanthomonas oryzae*, as reported by (57). **Geosmin** (F06) has been reported to act against *Aspergillus flavus* and *Aspergillus parasiticus* (35, 38), while **2-tetradecanone** (F64) has antibacterial effects on *Bacillus* species (53). Schulz and colleagues reported that **benzenecarbothioic acid, S-methyl ester** (F32) exhibited notable antifungal activity against *Aspergillus fumigatus*, *Botrytis cinerea*, and *Candida albicans*, as well as antibacterial activity against an *E. coli tolC* mutant (30). **Pyridine, 2,4,6-trimethyl** (F17) inhibited growth of *S. sclerotiorum* (56). Additionally, **2-tridecanone** (F53), **2-heptanone, 5-methyl** (F25) (21), **benzyl methyl ketone** (F76) (21) and **decane, 4-methyl-** (F61) (44) have been detected in extracts with antifungal activity, although their effects as pure compounds remain untested. Although **benzonitrile** (F14) is less toxic than many other compounds with a nitrile functional group, it serves as a precursor for various toxic and antimicrobial derivatives. Consistent with this, benzonitrile has been detected in antifungal extracts active against the phytopathogenic fungus *Moniliophthora roreri*. Interestingly, in contrast to its inhibitory properties, benzonitrile was found to stimulate the growth of *Pseudomonas fluorescens* (31–33).

Other bioactivities have been reported for three additional volatile compounds (VCs): the nitric oxide synthase inhibitor **3-penten-2-one** (F38) (22), the neurotransmitter, and cholinesterase inhibitor **phenol, 4-(2-aminoethyl)** (F67) (40–42) and **methyl benzoate** (F11), which has been shown to function as an insect attractant and insecticide, as well as to trigger induced systemic resistance in plants (24–26). In addition to this plant related compound, 7 other volatile compounds (F68, F79, F61, F47, F53, F43, F77) were directly or indirectly linked to the modulation of plant development and responses. Finally, **2-methylisoborneol**, which together with geosmin, modulates microbial predator-prey interactions by inhibiting bacterial grazing through chemical deterrence (37). Although the biological activity of pure **2-pentanone, 1-phenyl** (F46) has not been extensively studied in the literature, a study by Kuwahara et al. 2017 (47) highlighted its role in sexual signaling in the millipede *Nedyopus tambanus mangaesinus*.

The analysis of the variation of peak area values of the 36 VCs illustrates how the culture medium could influence the diversity of biological activities within a volatilome. The MHB and ISP1 media unambiguously best trigger the production of VCs, including some of those exhibiting multiple bioactivities including 2-undecanone (F43), 2-tridecanone (F53), 2-heptanone (F77), and DMTS (F58), which are found in categories with relatively high values. Interestingly, the compound with the highest overall peak area is methylthio-2-propanone (F19), a sulfur-containing compound produced most abundantly in ISP1, suggesting it may be a novel bioactive metabolite which functions remain to be discovered. Finally, VCs produced most abundantly in LB and TSB appear less frequently among the most abundant groups and generally show moderate peak area values, indicating that these media may be less favorable for significant volatile production. An exception is F65, significantly detected in TSB, though its function remains unknown.

### Inhibition by pure compounds

To identify the VCs produced by *S. scabiei* 87-22 that are responsible of the observed antifungal activities, we first selected DMTS (F58), 2-heptanone (F77), creosol (F06), and 2-phenylethyl acetate (F40), all previously reported for their antimicrobial properties. These four selected VCs exhibited antifungal activity against *A*. *solani*, *G*. *zeae*, and *P*. *restrictum* NS1, though to varying degrees. Additionally, 3-penten-2-one (F38) was tested due to its strong Log_2_ FC value (Table 1). Although this compound has never been previously characterized as an antimicrobial agent, our assays revealed potent antifungal activity against all tested fungi. This suggests that its known role as a nitric oxide synthase (NOS) inhibitor in mammalian cells may contribute to its antifungal properties. Among the tested compounds, 3-penten-2-one (F38) and DMTS (F58) completely inhibited all three fungal strains when pure VCs were diluted up to 10^2^, corresponding to approximately concentration values of 7.3 and 11.7 g/l, respectively. Additionally, 2-heptanone also demonstrated complete growth inhibition across all three strains but total inhibition was lost at a 10-fold dilution.

Creosol was previously demonstrated to have 100% antifungal activity against pathogenic fungi of the genera *Fusarium*, *Penicillium* and *Aspergillus* (*Fusarium oxysporum*, *Fusarium verticillioides*, *Penicillium brevicompactum*, *Penicillium expansum*, *Aspergillus flavus*, *Aspergillus fumigatus)* in agar dilution assays (39). However, when tested as a pure VC in bicameral Petri plates, its inhibitory effect against *A. solani*, *G. zeae*, and *Pe. restrictum* NS1 was limited, with no complete growth inhibition observed. Finally, 2-phenylethyl acetate reproducibly exhibited stronger antifungal activity at a 10-fold dilution compared to more concentrated conditions. This effect is particularly evident with *Pe. restrictum* NS1 (see Figure 6). This counterintuitive observation, known as the Eagle effect (or the paradoxical zone phenomenon), underscores the complexity of identifying bioactive compounds within a volatilome based solely on their relative peak areas.

**Figure 6.**
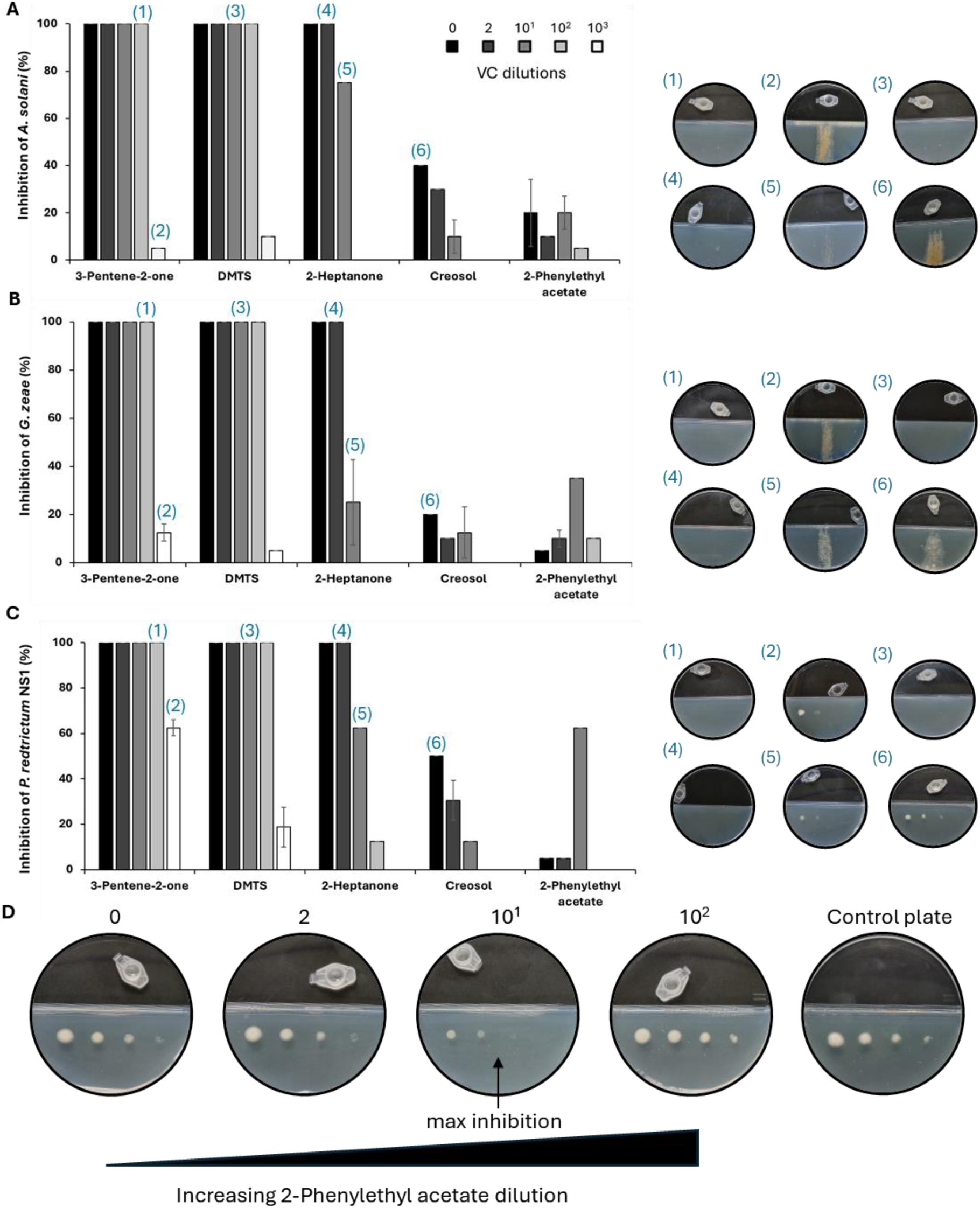
Antifungal activity of a selection of most abundant VCs produced by *S. scabiei*. Growth inhibition (%) of *A*. *solani* **(A),** *G*. *zeae* (**B**), and *Pe*. *restrictum* NS1 (**C**) by the different VCs tested as pure compounds. Dilutions of pure compounds from 0 (no dilution) to 10^3^ are color-coded. (**D**) Evidence of the Eagle effect observed for 2-pheniethyl acetate on *Pe. restrictum* NS1.

## Discussion

After showing that the nitrogen sources used as fertilizers impair growth of fungi by *S*. *scabiei* through siderophore-mediated iron sequestration (16), we now uncover a novel mechanism by which this species antagonizes other microorganisms. In this work, we explored by GC × GC-TOFMS the volatilome of *S. scabiei* 87-22 under five culture conditions known to induce the production of antifungal VCs. We identified 36 VCs unambiguously resulting from the metabolic activity of *S. scabiei* under at least one of the five tested conditions. Among the identified VCs, DMTS, 2-heptanone, creosol, 2-heptanone, 6-methyl, and 2-decanone, have been previously reported to inhibit fungi phylogenetically related to those tested in this study. Here we demonstrated that DMTS, creosol, and 2-heptanone, also inhibit the growth of the potato pathogen *Alternaria solani*, the cereal crop pathogen *Gibberella zeae*, and the non-pathogenic *Penicillium restrictum* NS1, expanding the known spectrum of fungal species sensitive to these VCs. Notably, DMTS has also shown significant efficacy against the oomycete *P. infestans*, the causative agent of late blight disease. Given the radically different severity and impact of late blight compared to common scab on plant viability, DMTS production upon colonization of potato plants by *S. scabiei* may not only positively influence its settlement in its environmental niche, but also mitigate the effects of much more virulent pathogens.

### Uncovering new bioactive compounds and unexpected behaviors

Our results also revealed the previously unreported antifungal activity of 3-penten-2-one. This compound, not previously recognized as an antimicrobial agent, exhibited strong antifungal activity against all tested fungi. Given its role as a nitric oxide synthase (NOS) inhibitor in mammalian cells, this newly identified activity warrants further investigation to understand its synthesis, perception, and broader ecological role in *S*. *scabiei*’s environment, including its interactions with neighboring microbes and host plants. Moreover, our study also uncovered unexpected behaviors of known volatiles, notably the paradoxical effect observed for 2-phenylethyl acetate exhibiting higher antifungal activity when diluted. This phenomenon known as the "Eagle effect" is characterized by the ability of specific strains to develop in the presence of high concentrations of an inhibitor while remaining vulnerable to lower concentrations. This effect has been reported in fungal species of *Aspergillus* and *Candida* treated with caspofungine or micafungine (68, 69) but no prior studies have documented this phenomenon in the fungal strains analyzed here.

In addition, *S. scabiei* 87-22 produces numerous other compounds known for their antimicrobial activity against a broader range of fungi and bacteria. However, further investigation is needed to determine their full antimicrobial potential as pure compounds. Overall, 12 volatile compounds from *S. scabiei* 87-22 have documented antimicrobial properties, highlighting its potential not only for niche competition but also for plant protection against more aggressive pathogens.

### Beyond antimicrobial activity: Plant-growth promotion and systemic resistance

Beyond compounds with antimicrobial activities, we identified VCs with activities promoting plant growth or plant-protecting activities while inducing systemic resistance. These findings challenge the conventional view of *S. scabiei* 87-22 solely as a pathogen, suggesting that its role in agroecosystems may be more complex than previously thought. Indeed, in most developing countries common scab is considered a cosmetic problem of minor importance, and our findings suggest that the volatile metabolites of *S. scabiei* 87-22 may enhance plant health in ways that have been largely overlooked. For example, 2-heptanone positively influenced the growth of *Arabidopsis thaliana*, while other alkyl ketones such as 2-decanone (34) and 2-tridecanone (46) have previously been identified in extracts that stimulate potato growth. Given that VCs trigger systemic signals within the plant and neighboring plants (70, 71), their impact extends beyond the sites of perception and exposed tissues, potentially contributing to plant growth promotion and protection at the whole plant level and its immediate community.

### Future directions and ecological implications

Obviously, it is essential to investigate whether these compounds are also emitted by *S. scabiei* 87-22 in its natural environment and how they function in host-pathogen dynamics and microbial communication and competition. Laboratory studies typically focus on the effects of individual compounds against a limited number of organisms, whereas in nature, volatile activity is influenced by a complex network of environmental factors. This variability emphasizes that the concentration, ratio, and synergistic interactions of volatiles determine their ultimate function within microbial communities. Therefore, in accordance with the principle *sola dosis facit venenum* (the dose makes the poison), determining the precise concentration and ratio of VCs needed to suppress pathogens while promoting plant growth would offer valuable insights with broad agricultural implications.

Finally, for some of the identified volatile compounds, no biological activity has been reported so far. These compounds deserve further investigation as pure substances to reveal their potential roles in shaping microbial rhizosphere, in sending chemical signals to host plants and their potential applications in plant protection and growth promotion. As traditional antimicrobial treatments become less effective, there is growing interest in exploring the diverse range of VCs produced by microorganisms like *Streptomyces*, particularly given their vast biosynthetic potential. This increasing interest is reflected in the rising number of publications addressing this topic. Estimating the exact number of VCs produced by *Streptomyces* species is challenging. However, a survey by Jones et al. (72) estimated that the volatilome of the *Streptomyces* genus comprises approximately 1,400 compounds spanning various chemical classes and biological activities. The identification of novel bioactive volatiles could offer sustainable alternatives for plant protection and green solutions for agriculture.

## Materials and methods

### Strain maintenance and cultivation media

Spore stocks of *S*. *scabiei* 87-22 (73) grown on IPM agar plates were prepared as in described in Practical Streptomyces Genetics (74) stored at -80°C or -20°C. *S*. *scabiei* 87-22 cultivation media used were (for 1 liter): International Streptomyces Project (ISP) media (N°1-7) prepared according to Shirling and Gottlieb (75); YM medium (Yeast extract (VWR^™^) 3 g, Bacto™ Malt Extract (Gibco™) 3 g, D(+)-Glucose monohydrate (Carl Roth^®^) 10 g, Bacteriological Peptone (Condalab) 5 g, and Bacto Agar (BD Difco™) 20 g); R2YE and SFM (Soy Flour Mannitol) media were prepared according to the protocol described in (74); IPM (Instant Potato Mash) medium consists of 50 g of Maggi Mousline (Nestlé^®^) powder and 12 g of Bacto Agar (BD Difco™) dissolved in tap water; TSA (Tryptic Soy Broth (MerckMilipore_®_) 30 g and Bacto Agar (BD Difco™) 20 g); MHA (Mueller-Hinton Broth (MerckMilipore_®_) 22 g and Bacto Agar (BD Difco™) 20 g); PDA (Potato Dextrose Broth (Sigma Aldrich^®^) 24 g/l and Bacteriological Agar (VWR^™^) 20 g); LB (Lysogeny Broth 25 g/l and Bacteriological Agar (VWR^™^) 20 g). V8 agar medium was prepared by dissolving 15 g of Bacto Agar (BD Difco™) and 3 g of calcium carbonate in 200 ml of V8^®^ vegetable juice, which was then mixed with 800 ml of distilled water; the pH was adjusted to 7.2.

Precultures of *Penicillium restrictum* NS1 (MUCL 58442), *Alternaria solani* (Ellis & Martin) Sorauer, *Gibberella zeae* (CIP collection strain), and *Fusarium culmorum* (MUCL42823) were prepared as follows: a 40% glycerol mycelium stock preserved at -80°C was used to inoculate PDA plates and incubated 3 days at 28°C; A V8 agar plate was inoculated with a mycelium plug of *Pythium ultimum* (DSM 62987). After 5 days at room temperature, new mycelium-rich plugs were taken from this plate and used to further assays.

### Antimicrobial assays

#### Cross-streak assays

The antimicrobial activity was first evaluated by cross-streak assays on solid media as previously described (76, 77). For *S*. *scabiei* 87-22, a 4 µl spore suspension (5.10^6^ spores/ml) was used to inoculate a single streak in the upper part of the Petri dish and plates were incubated at 28°C for 4 days. Mycelium suspension of *A. solani*, *G. zeae* and *F. culmorum* (OD_625_ at 0.1± 0.02 in water for mycelium scrapped from a 3-day culture at 28°C on PDA), was inoculated with a cotton swab as a single streak perpendicular to the band of *Streptomyces*. In anti-oomycete assays against *Pythium ultimum*, 7-mm plugs of *Py. ultimum* from 5 to 7-day-old cultures on solid V8 medium were placed into holes in the center of the plates, positioned 3 cm from the 4 days old *S. scabiei* streak. The plates were incubated at 28°C for 48 to 72 hours. The growth inhibition was measured by comparing the growth of tested microorganisms on the control plates with growth on plates containing *S. scabiei* 87-22. All experiments were performed in triplicate.

#### Assays for volatile compounds

Growth inhibition activity due to the production volatile compounds was assessed with two-compartment Petri dishes as described by Avalos et al. (5). One compartment is filled with the different media for *Streptomyces* growth and metabolite production, and the second compartment is filled with the PDA medium for fungal growth. *S. scabiei* 87-22 was inoculated in its compartments (a confluent lawn with 10 µl of a 10^7^ spores/ml suspension) 4 days at 28°C before. Petri dishes were sealed with parafilm to limit the produced volatile metabolites to escape. Mycelium of each fungal strain was scraped from a 3 days-old PDA preculture plate and suspended in sterile water to achieve an OD625_nm_ of 0.1 ± 0.02. Three dilution series of *Pe. restrictum* NS1 were prepared by diluting the suspension 10, 100, and 1000 times in sterile water; 5 μL of each dilution was spotted onto the second compartment, with the fourth spot being the undiluted mycelium suspension at OD625_nm_ = 0.1. For *A. solani* and *G. zeae*, the mycelium suspension at OD625_nm_ = 0.1 ± 0.02 was inoculated in form of streaks. Plates were incubated at 28°C for an additional 2 days to assess inhibition by the VCs produced by S*. scabiei* 87-22. Triplicate experiments were conducted for each condition.

#### Inhibitory effects of volatile compounds

The antifungal activity of five volatile compounds: 3-Penten-2-one (85%), Dimethyl trisulfide (98+%), 2-Heptanone (99%), creosol (99%) and 2-phenylethyl acetate (98%) (purchased from Thermo Fisher Scientific®) was assessed in bicameral Petri dishes (SARSTEDT). In the experimental setup, one compartment contained PDA for the growth of fungal strains (*A. solani, G. zeae*, and *Pe. restrictum* NS1), while the adjacent compartment contained a small container holding 100 µl of tested compound. The preparation and inoculation procedures for the fungal strains were performed as described above. Plates were sealed with parafilm and incubated individually in hermetically closed containers at 28°C for an additional 2 days to assess the inhibition caused by the volatile compounds. Experiments were repeated twice.

#### Sampling

Thirty milliliters of each broth culture (ISP1, ISP6, LB, TSB, MHB) were inoculated with 10^7^ spores of *S. scabiei* 87-22. Both inoculated and non-inoculated duplicate cultures were incubated for 3 days at 28°C with agitation at 180 RPM. After incubation, the cultures were centrifuged at 4°C for 15 minutes at 4000 RPM. The supernatants were then filtered through a 0.22 µm filter. For analysis of culture media and bacterial samples, 5 ml aliquots were transferred into 20 ml headspace vials. From each of the duplicate cultures, three 5 ml aliquots were taken from the inoculated condition to generate three technical replicates, and two 5 ml aliquots were taken from the non-inoculated condition to produce two technical replicates.

These vials were stored at -20 °C and removed from the fridge on the same day as they were analyzed. The samples were first incubated at 40 °C for 10 minutes with an agitator speed of 250 rpm. Then, the headspace of each sample was extracted for 35 minutes using a DVB/CAR/PDMS SPME fiber (Supeclo®, Bellefonte, Pas, USA), which was selected for its broad-range adsorption capabilities. This fiber combines divinylbenzene (DVB), carboxen (CAR), and polydimethylsiloxane (PDMS), making it particularly suitable for capturing a wide variety of volatile and semi-volatile compounds with different polarities and molecular weight. The SPME fiber was first conditioned according to the supplier’s instructions before use. During measurement, the fiber was pre-conditioned at 270°C for 10 minutes and post-conditioned at 270°C for 15 minutes to ensure that all adsorbed compounds were desorbed. The bacterial and media samples were injected in a randomized order to prevent any patterns that could bias the results. This approach helps ensure the data fairly and accurately represents the different conditions, leading to more reliable conclusions. Quality control (QC) samples were included throughout the analysis sequence to ensure the accuracy and consistency of the analytical process. 1 µl of a QC 37 mixture (In-house QC mixture) was introduced into 20 ml headspace vials and analyzed alongside the samples. To establish a reliable baseline for the QC Chart, ten replicates of the QC 37 mixture were injected at the beginning of the study. This allowed for the assessment of instrument stability and method reproducibility over time. To determine linear retention indices, 1 µl of a 5 ppm n-alkane mix (Hexane (C_6_) - Eicosane (C_20_)) was injected using the same conditions as for the sample analyses.

#### GC × GC-TOFMS Method

The bacterial sample analysis was performed with Pegasus BT 4D GC × GC-TOFMS system with a thermal quad jet modulator (LECO Corp., St. Joseph, MI, USA) equipped with a CTC PAL RSI autosampler (CTC analytics, Switzerland). To achieve an adequate separation, the column set consisted of a normal-phase (non-polar × mid-polar) composed of first dimension (^1^D) Rxi-5 Sil MS (30 m × 0.25 mm × 0.25 μm) and second dimension (^2^D) Rxi-17 Sil MS (2 m × 0.25 × 0.25 μm). This column combination offers complementary selectivity, the non-polar ^1^D column separates compounds mainly based on volatility, while the mid-polar ^2^D column introduces additional separation based on polarity. This orthogonality enhances overall resolution, allowing for improved characterization of the complex bacterial profile.

Headspace samples were desorbed from SPME fiber in the GC injector at 250 °C for 10 min. The main GC oven was set at 40 °C for 1 min, then increased to 200 °C with a ramp of 5 °C/min, and finally increased to 270 °C (held for 2 min) with a ramp of 20 °C/min, for a total run of 38.30 min. The secondary oven temperature offset was 15°C, and the modulation period (PM) was set at 2 s. The carrier gas used was Helium at 1.4 ml/min, the system was equipped with a split/spitless injector set on a 1: 10 ratio. The Mass spectrometer (MS) was a time-of-flight (TOF) instrument. The MS parameters were as follow: source temperature at 250 °C, MS transfer line temperature at 250 °C, the ionization mode was electron ionization (EI) at 70 eV, MS was operated in a full scan mode for *m/z* 35-450, and the acquisition rate was 200 spectra/ seconds.

### Data processing

Raw data acquisition and pre-processing were performed using ChromaTOF® software (version 4.72, LECO Corp., St. Joseph, MI, USA). Putative identification of analytes was conducted by matching mass spectra against the NIST library. For comparative analysis of bacterial and media chromatograms, ChromaTOF® Tile (version 1.01, LECO Corp.) was used. This software is specifically designed to handle complex GC × GC datasets by dividing each chromatogram into a grid of small sections known as “tiles.” This approach simplifies data and facilitates direct comparison across samples. To ensure consistency, the software also aligns retention times between chromatograms, correcting for slight chromatographic shifts.

In this study, a tile size of 8 modulations in the first dimension (^1^D) and 150 spectra in the second dimension (^2^D) was used, with a signal-to-noise (S/N) threshold of 50. ChromaTOF® Tile applies a Tile-Based Fisher Ratio (TBFR) algorithm to statistically evaluate differences between predefined sample groups. In this case, inoculated versus non-inoculated media. The Fisher ratio (F-ratio) measures a compound’s ability to discriminate between groups, with higher values indicating stronger differentiation. F-ratios in this analysis ranged from 12 to 537. For downstream analysis, only compounds with an F-ratio greater than 50 were retained, resulting in the selection of 79 discriminative compounds.

The chemometric tests including PCA and volcano plot were performed using MetaboAnalyst 5.0 (Xia Lab, McGill University, Montréal, QC, Canada). For the chemometric analyses, the areas of the detected compounds were used. Prior to analysis, the data underwent median normalization (to reduce signal intensity discrepancies), logarithmic transformation (to reduce the impact of data skewness), and autoscaling (to center and scale the variables).

## Data availability

GC×GC-TOF-MS VCs Dataset are available at https://zenodo.org/records/15265340

## Acknowledgments

The work of N.S. was supported by a FRIA grant from the "Fonds de la Recherche Scientifique" (FRIA 1.E.116.21), and a FNRS grant "Crédit de recherche" (grant CDR/OL J.0158.21) to S.R. Support was also provided to D.B. by the graduate school for research XL–Chem (ANR-18-EURE-0020 XL CHEM), and to PH.S. by Léon Fredericq foundation.

## Supplementary files

**Figure S1:**
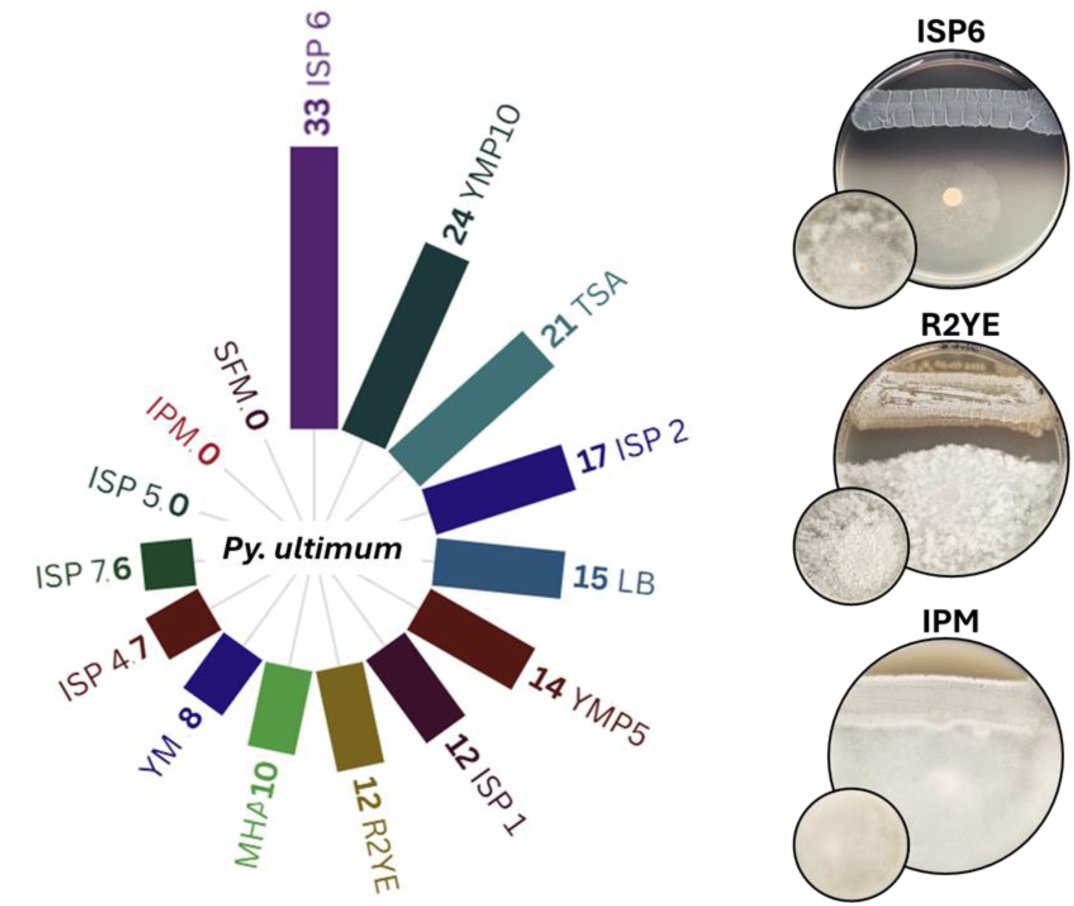
Semi-quantitative evaluation of antimicrobial activities of *S*. *scabiei* 87-22 against oomycete *Pythium ultimum* on 15 growth media. A score of 100 means full growth inhibition of the tested microorganisms, while a score of 0 indicates no growth inhibition. Insets in the bottom left corner are control plates which display the growth of tested microorganisms without *S. scabiei.* Chart created using the Flourish.studio (https://flourish.studio).

**Figure S2.**
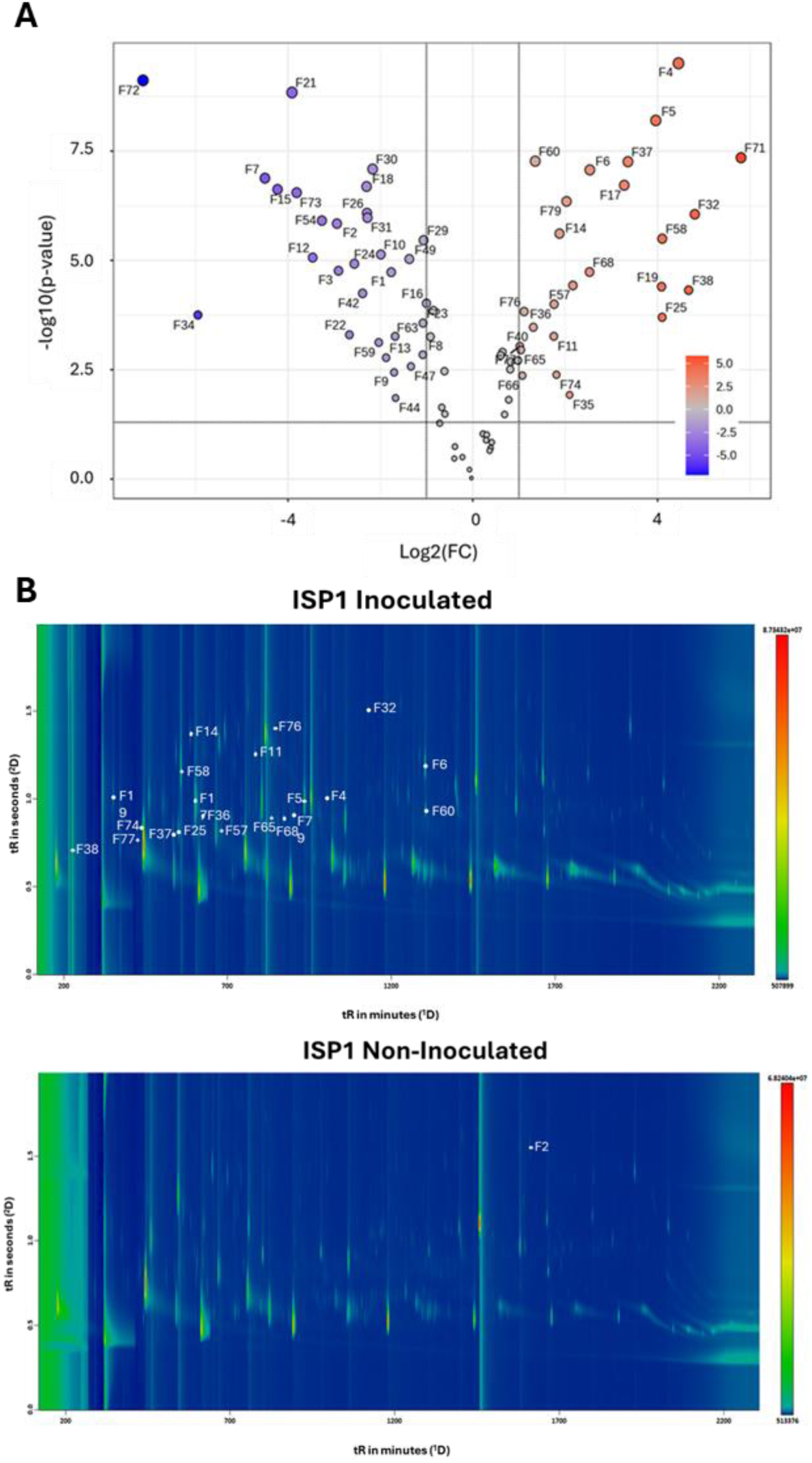
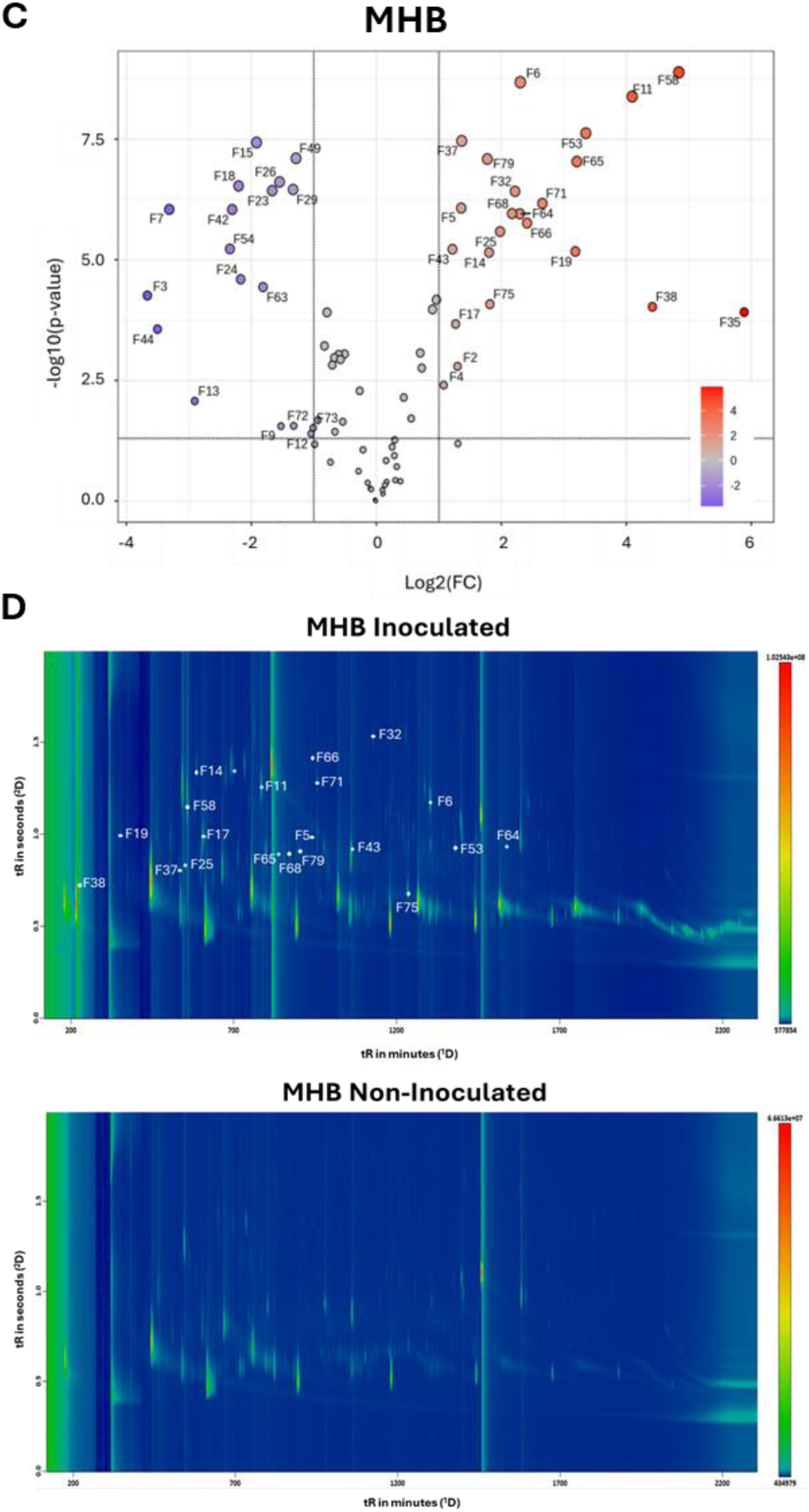
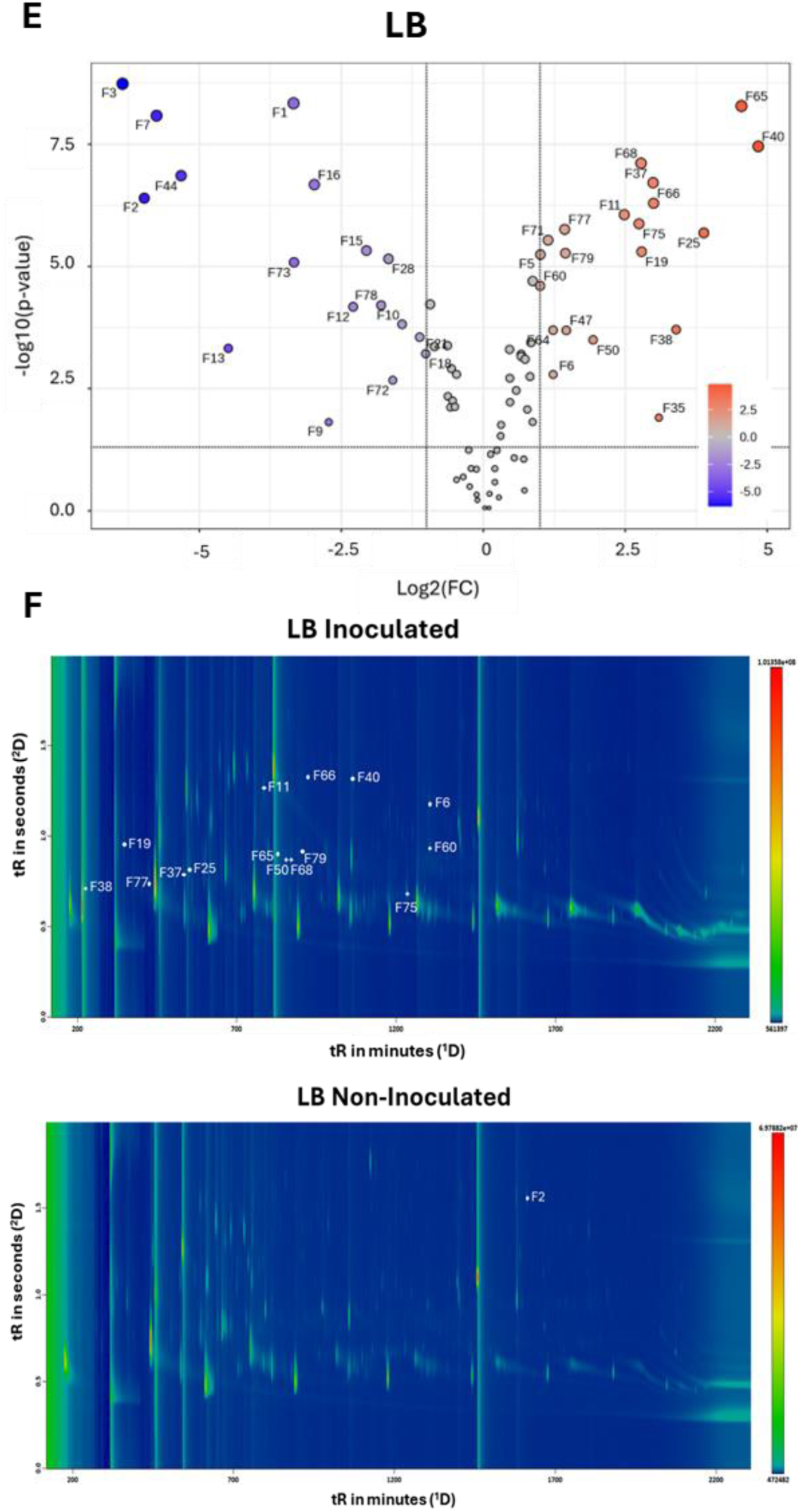
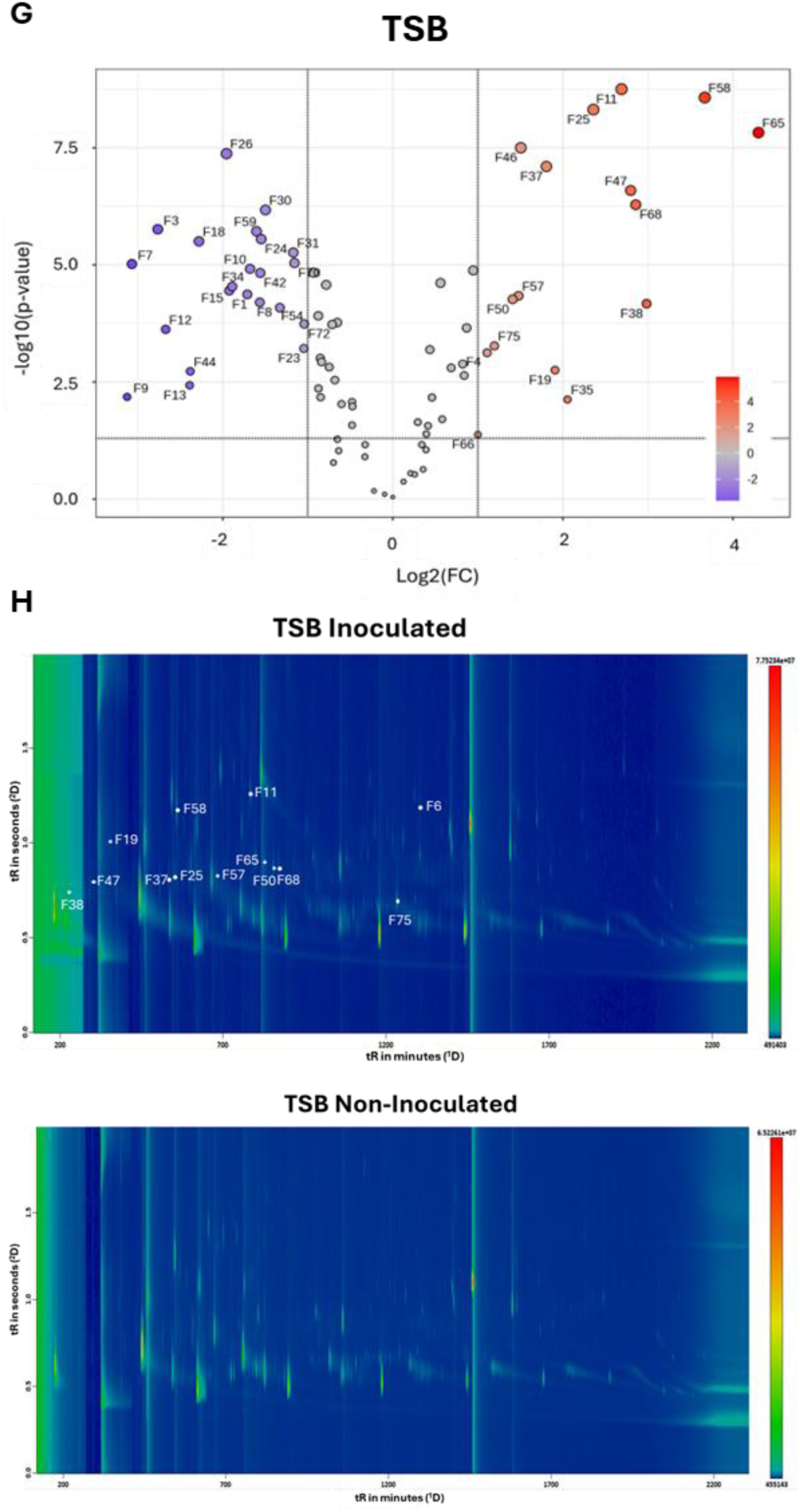
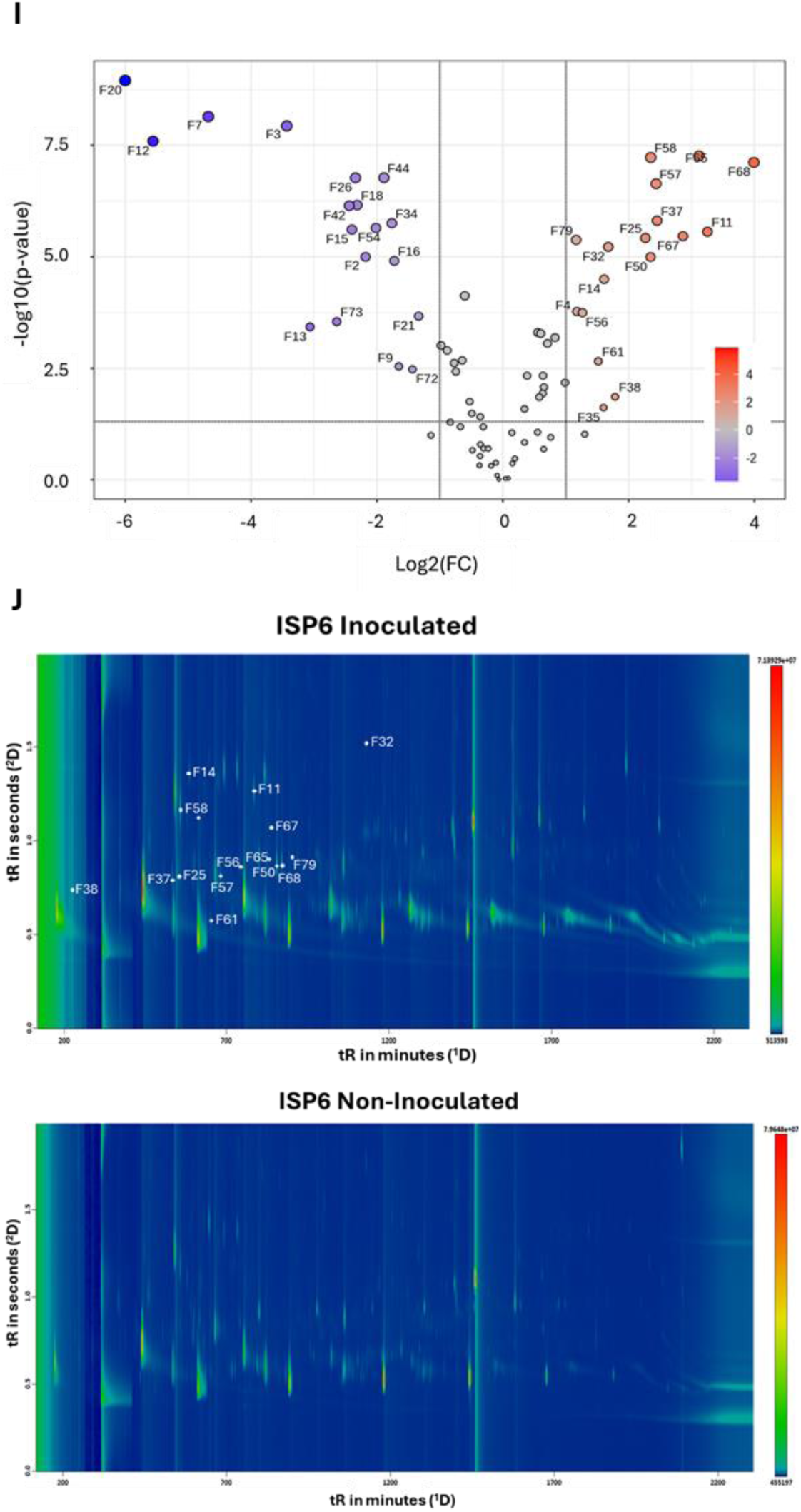
Volcano plots and GC × GC -TOF MS chromatograms in ISP1 (A, B), in MHB (C, D), in LB (E, F), in TSB (G, H) and in ISP6 (I, J). Upper panels: volcano plots of VCs identified by GC × GC-TOFMS across five media. The X-axis shows the Log₂ FC in VC intensity between medium non-inoculated (control) and inoculated by *S. scabiei* 87-22. The Y-axis displays the -log₁₀ (p-value), signifying the statistical significance of these changes (p-value ≤ 0.05 or -log_10_ p-value ≥ 1.3). The volcano plots are divided into three zones: 1) where the VCs do not result from the metabolic activity of *S. scabiei* 87-22 (Log_2_ FC < -1, blue circle); 2) where the log₂ FC did not significantly vary (-1 < Log_2_ FC < 1, grey circle), and 3) where result from the metabolic activity of *S. scabiei* 87-22 (Log_2_ FC ≥ 1, red circle). Middle and lower panels: chromatograms of inoculated and non-inoculated media are represented in lower panels. Each spot in the chromatogram corresponds to a VC separated by GC × GC: the first dimension (^1^D), using a non-polar column, separates compounds primarily according to their boiling points, while the second dimension (^2^D), using a mid-polar column, separates them based on their polarity. The area intensity of each compound is visualized using a color gradient, from light blue (lower intensities) to red (higher intensities), the dark blue color represents the background noise.

**Supplementary Table S1.** MSI-based identification levels and spectrometric data of volatile compounds with significant Log2 FC ≥ 1.

| Feature number | Compound name | RI | RI Lib | Δ RI | Similarity | Reverse | MSI<sup>1</sup> Level |
| --- | --- | --- | --- | --- | --- | --- | --- |
| F02 | 2-p-Tolylamino-cyclopent-1-enecarbonitrile | 1623.9 | / | / | 703 | 717 | 3 |
| F04 | Acetamide, N-(2,6-dimethylphenyl)- | 1222.9 | / | / | 785 | 802 | 3 |
| F05 | 2-Methylisoborneol | 1183.9 | 1197 | 13.1 | 724 | 728 | 3 |
| F06 | Geosmin | 1408.1 | 1431 | 22.9 | 838 | 880 | 3 |
| F11 | Benzoic acid, methyl ester | 1095.6 | 1096 | 1.6 | 940 | 942 | 2 |
| F14 | Benzonitrile | 985.1 | 985 | 0.1 | 967 | 967 | 2 |
| F17 | Pyridine, 2,4,6-trimethyl- | 993.1 | 991 | 2.1 | 959 | 962 | 2 |
| F19 | Methylthio-2-propanone | 850.6 | 863 | 12.4 | 879 | 879 | 2 |
| F25 | 2-Heptanone, 5-methyl- | 965.5 | 971 | 5.5 | 876 | 924 | 2 |
| F32 | Benzenecarbothioic acid, S-methyl ester | 1297.6 | 1299 | 1.4 | 939 | 939 | 2 |
| F35 | Dimethyl selenodisulfide | 1048.9 | 1047 | 1.9 | 705 | 750 | 3 |
| F36 | Benzenamine, N-(1-methylethyl) | 1007.8 | / | / | 741 | 741 | 3 |
| F37 | 2-Heptanone, 6-methyl- | 955.2 | 956 | 0.8 | 843 | 881 | 2 |
| F38 | 3-Penten-2-one | 777 | 735 | 42 | 930 | 943 | 1 |
| F40 | Acetic acid, 2-phenylethyl ester | 1259 | 1258 | 1 | 918 | 930 | 1 |
| F43 | 2-Undecanone | 1259 | 1294 | 35 | 848 | 854 | 3 |
| F46 | 2-Pentanone, 1-phenyl | 1312 | 1327 | 15 | 678 | 813 | 2 |
| F47 | 2-Hydroxy-3-pentanone | 821.8 | 821 | 0.8 | 826 | 882 | 2 |
| F50 | 4-Nonanone, 8-Methyl- | 1135.6 | / | / | 750 | 825 | 3 |
| F53 | 2-Tridecanone | 1460.8 | 1497 | 36.2 | 856 | 856 | 3 |
| F56 | 4-Octanone, 7-methyl- | 1073.3 | / | / | 826 | 866 | 3 |
| F57 | 4-Nonanone | 1038.9 | 1030 | 8.9 | 788 | 877 | 2 |
| F58 | Dimethyl trisulfide | 969 | 970 | 1 | 917 | 917 | 1 |
| F60 | Dodecanal | 1409.5 | 1409 | 0.5 | 916 | 930 | 2 |
| F61 | Decane, 4-methyl- | 1022.2 | 1060 | 37.8 | 857 | 876 | 3 |
| F64 | 2-Tetradecanone | 1569 | 1597 | 28 | 849 | 849 | 3 |
| F65 | 2,2-Dimethylheptane-3,5-dione | 1119.5 | / | / | 829 | 829 | 3 |
| F66 | Benzeneethanol, Beta-methyl- | 1174.7 | 1179 | 4.3 | 832 | 832 | 2 |
| F67 | Phenol, 4-(2-aminoethyl) | 1125.3 | / | / | 673 | 695 | 4 |
| F68 | 4-Decanone | 1144.8 | 1137 | 7.8 | 744 | 803 | 2 |
| F71 | Creosol | 1194.3 | 1193 | 1.3 | 912 | 912 | 1 |
| F74 | 3-Hexen-2-one, 5-methyl- | 900 | / | / | 887 | 913 | 3 |
| F75 | 2,5-Dimethylhexane-2,5-dihydroperoxide | 1364.6 | 1367 | 2.4 | 734 | 753 | 3 |
| F76 | Benzyl methyl ketone | 1129.9 | 1110 | 10.9 | 895 | 895 | 2 |
| F77 | 2-Heptanone | 893.1 | 891 | 2.1 | 924 | 941 | 1 |
| F79 | 2-Decanone | 1163.2 | 1193 | 29.8 | 784 | 803 | 3 |

<sup>1</sup> MSI: Based on The Metabolomics Standards Initiative (MSI), four identification levels were established, from Level 1 (highest confidence) to Level 4 (lowest), using criteria adapted for GC × GC data. These include the <sup>1</sup>D ΔRT, which is the difference in first-dimension retention time between the reference standard and the compound from the sample; the RSI (Reverse Similarity Index), which compares the compound's mass spectrum to a library spectrum; and the ΔRI, the difference between the experimental and library retention indices.
Level 1 : Identified Compounds
- Reliably identified using a commercial standard.
- Criteria: ΔRT (in-house) < 20 s, RSI > 800, and ΔRI < 20.
Level 2: Putatively Annotated Compounds
- No standard used.
- Identification based on spectral similarity with commercial libraries (MAINLIB, Replib, Nist\_RI).
- Criteria: RSI > 800 and ΔRI < 20.
Level 3: Putatively Characterized Compound Classes
- Assigned to a known chemical class based on spectral similarity with compounds from that class.

